# Regulation of dopamine release by tonic activity patterns in the striatal brain slice

**DOI:** 10.1101/2024.05.22.595411

**Authors:** Siham Boumhaouad, Emily A Makowicz, Sejoon Choi, Nezha Bouhaddou, Jihane Balla, Khalid Taghzouti, David Sulzer, Eugene V. Mosharov

**Affiliations:** Departments of Psychiatry and Neurology, Division of Molecular Therapeutics, New York State Psychiatric Institute, Columbia University Medical Center, New York, NY, USA; Physiology and Physiopathology Team, Genomics of Human Pathologies Research Center, Faculty of Sciences, Mohammed V University in Rabat, Rabat, Morocco

**Keywords:** dopamine, in vivo DA release, fast-scan cyclic voltammetry, striatal slice, D2 receptors, nicotinic acetylcholine receptor

## Abstract

Voluntary movement, motivation, and reinforcement learning depend on the activity of ventral midbrain neurons that extend axons to release dopamine (DA) in the striatum. These neurons exhibit two patterns of action potential activity: a low-frequency tonic activity that is intrinsically generated and superimposed high-frequency phasic bursts that are driven by synaptic inputs. *Ex vivo* acute striatal brain preparations are widely employed to study the regulation of evoked DA release but exhibit very different DA release kinetics than *in vivo* recordings. To investigate the relationship between phasic and tonic neuronal activity, we stimulated the slice in patterns intended to mimic tonic activity, which were interrupted by a series of burst stimuli. Conditioning the striatal slice with low-frequency activity altered DA release triggered by high-frequency bursts and produced kinetic parameters that resemble those *in vivo*. In the absence of applied tonic activity, nicotinic acetylcholine receptor and D2 dopamine receptor antagonists had no significant effect on neurotransmitter release driven by repeated burst activity in the striatal brain slice. In contrast, in tonically stimulated slices, D2 receptor blockade decreased the amount of DA released during a single burst and facilitated DA release in subsequent bursts. This experimental system provides a means to reconcile the difference in the kinetics of DA release *ex vivo* and *in vivo* and provides a novel approach to more accurately emulate pre- and post-synaptic mechanisms that control axonal DA release *in vivo*.

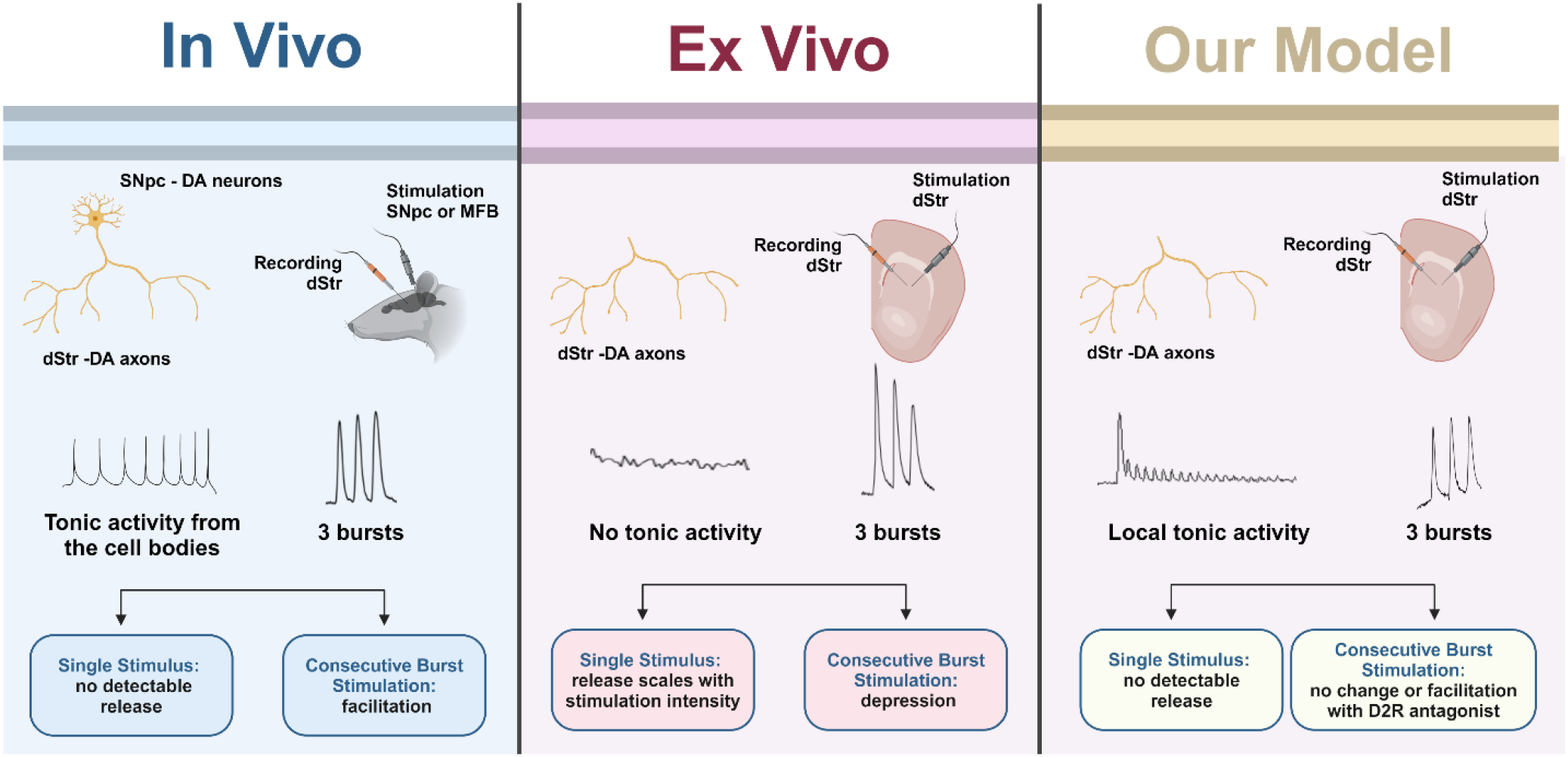

## 1. INTRODUCTION

Basal ganglia dopaminergic neurons located in the substantia nigra pars compacta (SNpc) and ventral tegmental area of the midbrain send axonal projections to the striatum, where the release of dopamine (DA) plays pivotal roles in reinforcement learning, motivation, and motor control ^1–3^. *In vivo*, these neurons predominantly exhibit two firing patterns: a tonic self-autonomous single-spike activity at 1-10 Hz ^4^, and phasic burst activity at 13-20 Hz in anesthetized rodents and up to 80 Hz in awake animals or humans ^5^ that is driven by synaptic inputs. Tonic and phasic firing of DA neurons have been suggested to play distinct roles in animal behaviors: tonic neurotransmitter release is thought to be required for motivation and motor control, whereas phasic activity in response to environmental cues encodes reward prediction error and is important for learning ^6,7^. Both tonic and phasic activity are regulated by pharmacological agents, including antipsychotic drugs, psychostimulants, and drugs of abuse including opiates, nicotine, and alcohol. Dysregulation of firing patterns is associated with neurological disorders including Parkinson’s disease, addiction, and schizophrenia ^8^.

Many studies have analyzed the kinetics of DA exocytosis in acute striatal slices, providing fundamental insights into the regulation of evoked neurotransmitter release by drugs and local synaptic circuitry and demonstrating regional variations in DA release probability and short-term plasticity within the striatum ^9–13^. These *ex vivo* studies have shown that striatal DA release is modulated by many mechanisms, including changes in vesicular DA content, inhibition of the dopaminergic axons by D2 receptors (D2R) ^14–19^, control by locally released acetylcholine (ACh) ^12,20^, and regulation of fusion pore kinetics ^21^. Additionally, the dynamic activity-dependent equilibrium between different pools of synaptic vesicles within the release sites – the readily releasable pool (RRP), the recycling pool and the reserve pool – is an important factor in the regulation of DA vesicle release probability and the refilling of synaptic vesicles with neurotransmitter ^21–25^. Due to these mechanisms, phasic DA release has been suggested to inversely depend on the frequency of tonic activity, due to a larger readily releasable pool of synaptic vesicles and lower D2R-mediated autoinhibition in inactive dopaminergic axons ^6,26^, although experimental evidence for this hypothesis is lacking.

While the acute striatal brain slice provides strong experimental advantages due to the ease of pharmacological or optogenetic manipulations of the local circuitry, limitations of this model system are highlighted by very different DA release kinetics *in vivo* ^22,27^. Typically, kinetic studies of electrochemical recordings of evoked DA release *in vivo* are performed in anesthetized animals with the recording electrode implanted in the striatum and stimulation electrode in the ventral midbrain or the medial forebrain bundle. In contrast to slice recordings, where a single electric pulse evokes large amounts of DA release ^18,28^, a single stimulus is typically insufficient to evoke detectable release of DA *in vivo* ^22,29^. Moreover, consecutive phasic bursts in striatal slices produce a profound release depression that contrasts with a facilitation of DA release observed *in vivo* ^22^.

A notable difference between the *in vivo* and *ex vivo* recordings is that tonic activity is present *in vivo* but absent in coronal striatal slices that lack the dopaminergic neuron cell bodies that generate tonic activity. To investigate whether tonic activity underlies the difference between *ex vivo* and *in vivo* kinetics of DA release and plasticity, we applied local electrical stimulation to the coronal slice of mouse dorsal striatum at various tonic frequencies. The tonic conditioning of the slice was followed by a series of high frequency bursts to determine changes in release facilitation or depression. Following characterization of these kinetics, we investigated the effect of nicotinic ACh receptor and D2 dopamine receptor antagonists on slice burst kinetics in the presence and absence of tonic activity. Our results indicate that superimposition of phasic firing patterns on tonic activity provides a means to use an *ex vivo* preparation for an improved analysis of the mechanisms that govern the release of monoamine neurotransmitters *in vivo*.

## 2. RESULTS

### 2.1. Effect of tonic stimulation on burst DA release

To verify previously reported differences between *in vivo* and *ex vivo* striatal recordings, we compared evoked DA release in anesthetized mice and in acute striatal slices after a single (1p) stimulus followed by three high-frequency bursts of 30 pulses at 50 Hz (30p; **Figure 1A**). *In vivo*, electrical stimuli were applied to the SNpc, while DA release was recorded in the dorsal striatum with fast scan cyclic voltammetry (FSCV). DA release in the striatal slice was stimulated locally in the dorsal striatum.

**Figure 1:**
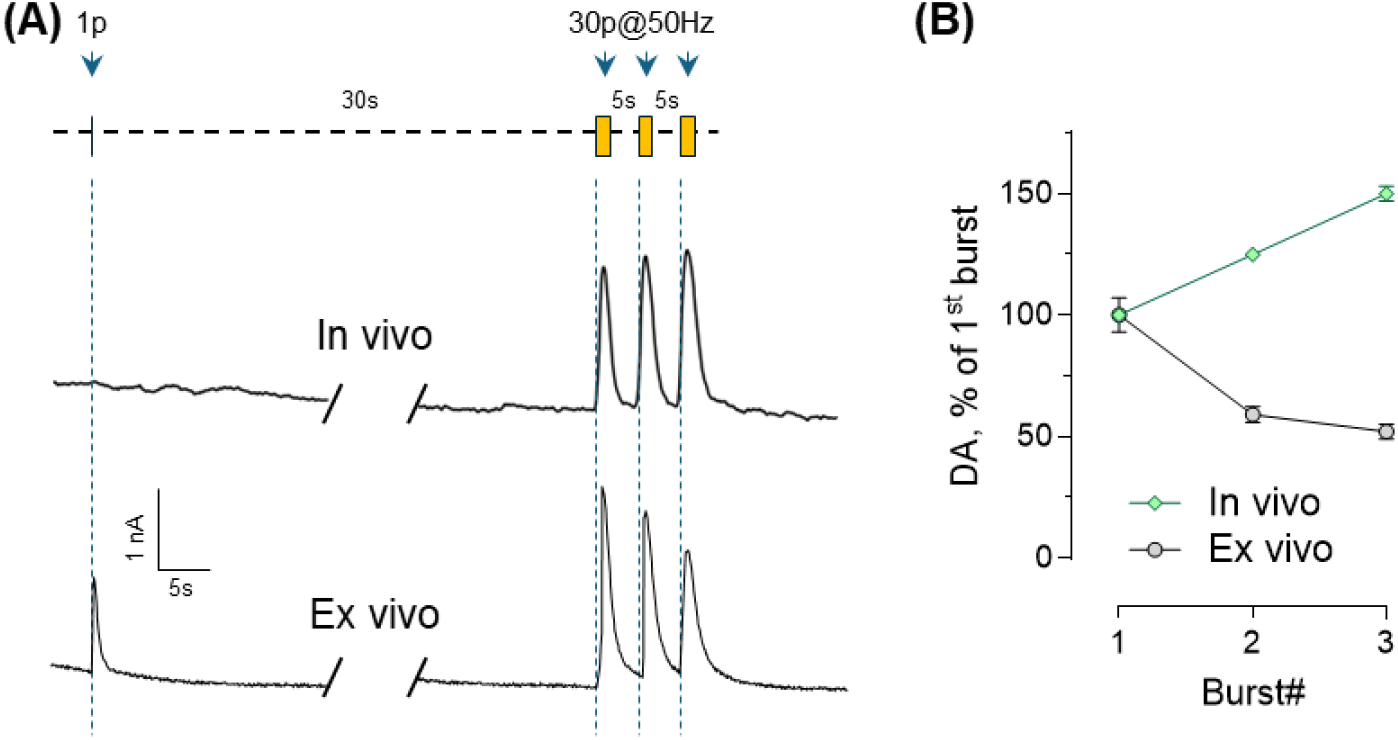
Comparison of DA release kinetics following single pulse and burst stimuli *in vivo* and *ex vivo*. Recordings from anesthetized mice *in vivo* were performed with FSCV electrode placed in the dorsal striatum and electrical stimulation in the SNpc. *Ex vivo* recordings were done in acute coronal dorsal striatum slices with local electrical stimulation. **(A)** Stimulation paradigm (top) and representative FSCV traces from *in vivo* (middle) and *ex vivo* (bottom) recordings. **(B)** Changes in the amplitude of DA peaks amplitude during a series of burst stimuli *in vivo* and in a slice. Curves are significantly different with p<0.0001 by 2-way ANOVA (n= 12 *in vivo* and 46 slice recordings).

As expected, *in vivo* DA release was below the detection limit with a single pulse stimulation and was only detectable after a burst of stimuli (**Figure 1A**). In contrast, 1p stimulus produced a large FSCV signal in the slice, with burst stimulation further increasing the amount of released neurotransmitter. Importantly, facilitation of DA release during consecutive stimulation bursts was observed *in vivo* while a depression of release occurred *ex vivo* (**Figure 1B**).

To investigate whether the lack of tonic activity in the slice is responsible for the differences in DA release kinetics *in vivo* and *ex vivo*, we stimulated slices at frequencies ranging from 0.5 to 5 Hz that were intended to emulate pacemaking activity of DA neurons (**Figure 2A**). We note that synaptic vesicle fusion in DA axons is expected to have a higher release probability with local electrical stimulation than with action potentials generated distally in the cell bodies, due to current loss along lightly myelinated dopaminergic axons ^30,31^. Introduction of artificial tonic activity in the striatal slice rapidly decreased DA release in individual pulses to undetectable levels, with more rapid depletion at higher stimulation frequencies (**Figure 2B-C**). No evoked DA peaks were observed at any tonic frequency stimulus by the end of a 5 minute train application.

**Figure 2:**
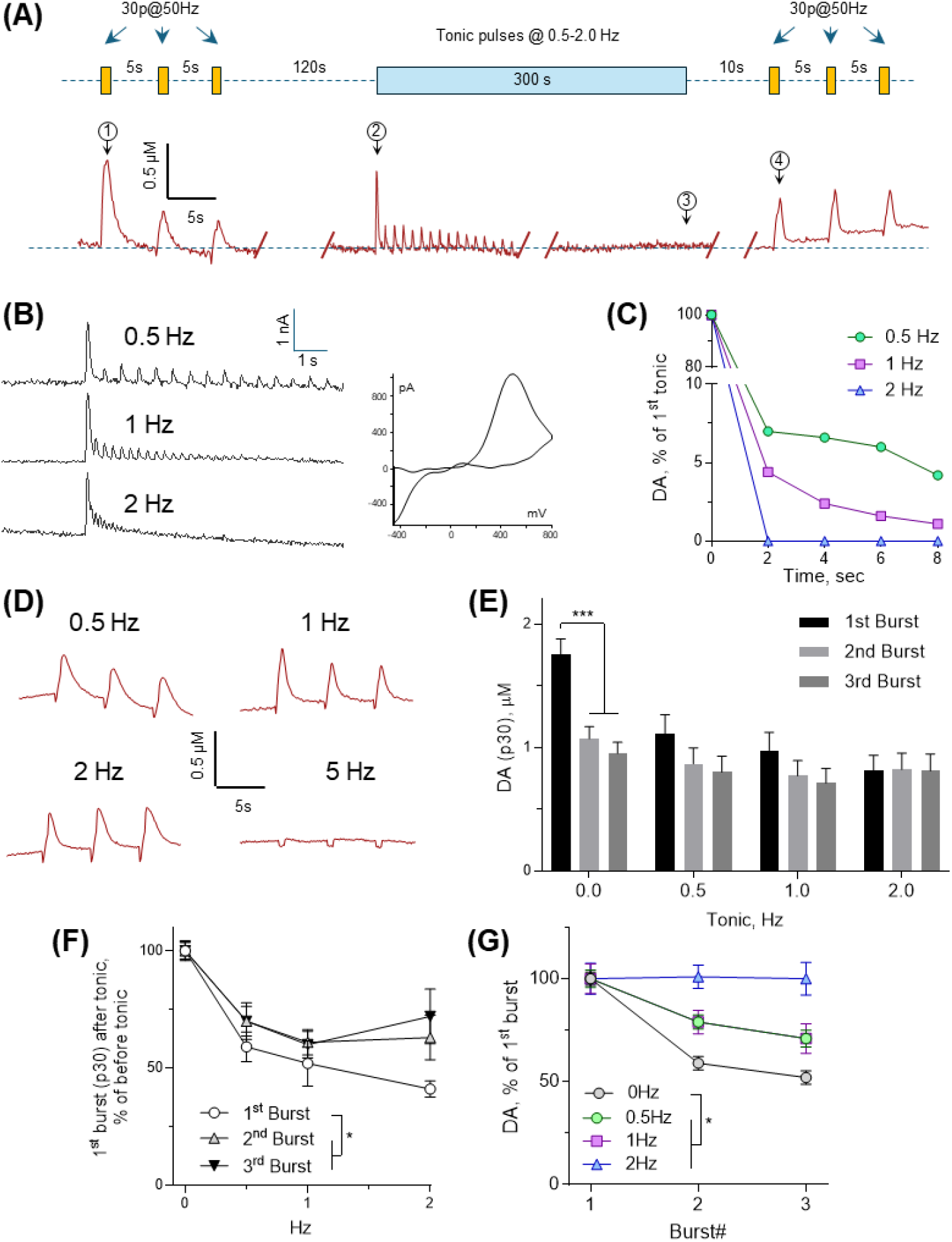
Effect of tonic frequency on the amplitude of evoked striatal dopamine release. **(A**) Stimulus paradigm used in each recording consisted of three phasic bursts (30 pulses at 50 Hz), followed by tonic stimulation at 0.5-5 Hz for 5 min, and then three more phasic bursts. Representative FSCV traces are shown below. Encircled numbers denote DA release following (1) 30p burst without tonic activity, (2) 1p stimulus without tonic activity, (3) 1p stimulus after tonic activity, and (4) 30p burst after tonic activity. **(B)** Representative FSCV traces of DA release peaks during the first 10 sec of tonic activity show frequency-dependent depression of DA release, followed by a complete cessation of neurotransmission. No DA release was detectable by the end of the 5 min tonic stimulation. Representative voltammogram of DA oxidation/reduction after the 1^st^ stimulus is shown on the right. **(C)** Average amplitudes of DA release peaks during the 1^st^ 10 sec of tonic activity. **(D)** Representative FSCV traces of burst stimulation-dependent DA release after 5 min of tonic slice conditioning. No burst DA release was detected after 5 Hz tonic stimulation. **(E)** Average amplitudes of DA release peaks following a series of stimuli bursts (***- p< 0.001 by one-way ANOVA; n= 46 0 Hz, 12 0.5 Hz, 12 1 Hz and 7 2 Hz). **(F)** Effect of preceding tonic slice stimulation on DA release evoked by phasic bursts 1-3. * - p<0.05 by two-way ANOVA. **(G)** Progressive relief from depression during a series of phasic bursts with increased tonic frequency. * - p<0.05 between curves by two-way ANOVA.

Tonic stimulation was followed by three bursts, each consisting of 30 pulses at 50 Hz, to mimic phasic activity - identical stimulus series was applied before the start of the tonic activity as an internal control for the effect of tonic stimuli (**Figure 2A**). An increase in tonic frequency from 0.5 to 2 Hz produced progressive reduction in DA release following a single phasic burst (**Figure 2E-F**), while no release was detected at 5 Hz, likely due to the depletion of DA stored in synaptic vesicles (**Figure 2D**). Interestingly, neurotransmitter release during the 2^nd^ and 3^rd^ high-frequency bursts was much less affected by preceding tonic stimulation (**Figure 2E-F**). This resulted in significantly attenuated DA release depression during a series of burst stimuli in the presence of tonic stimulation (**Figure 2G**), suggesting that depression phenomenon *ex vivo* is dependent on the lack of tonic activity.

### 2.2. Control of DA release by nAChR

Cholinergic interneurons (ChI) constitute only 2%–5% of striatal neurons, but establish an extensive network of axons that provides profound control over striatal DA neurotransmission via pre- and postsynaptic cholinergic receptors ^32–35^. Importantly, in contrast to the distal stimulation *in vivo*, local striatal slice stimulation *ex vivo* includes concurrent activation of local ACh release, leading to nAChR activation and desensitization. Similar to previously published results ^35–37^, application of the nAChR antagonist dihydro-β-erythroidine hydrobromide (DhβE, 1 μM) decreased DA release after a single stimulus but increased it after burst stimuli, thus significantly enhancing burst-to-single pulse ratio (**Figure 3A-C**). However, inhibition of nAChR in the presence of artificial tonic activity exerted no additional effect on neurotransmitter release evoked by a single burst (**Figure 3D**) or a series of bursts (**Figure 3E**), supporting the idea that nAChRs are desensitized and play a limited role in DA release modulation in the presence of artificial tonic activity.

**Figure 3:**
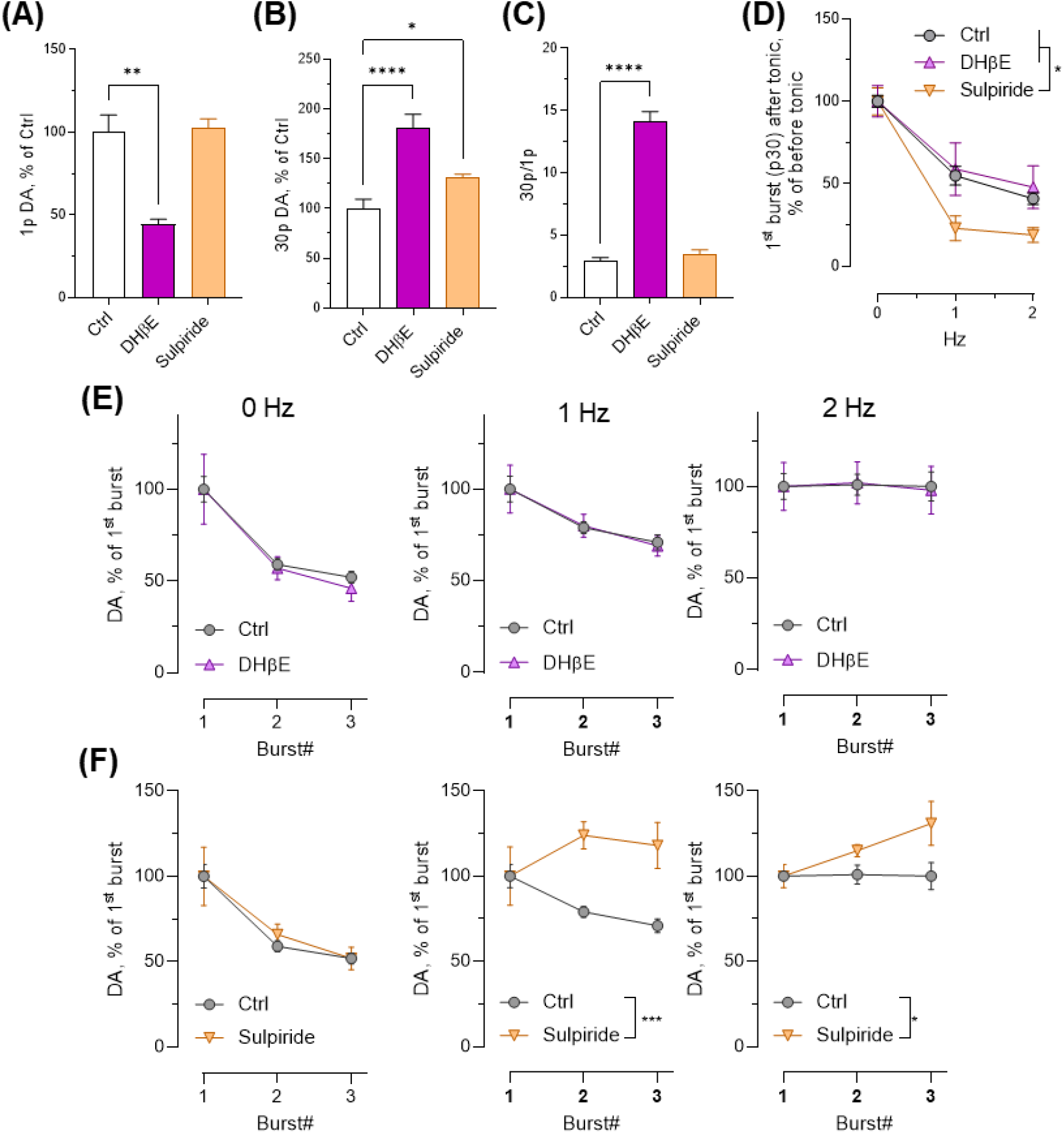
Modulation of evoked striatal DA release by nAChR and D2 antagonists. **(A-C)** Effect of nAChR antagonist DHβE (1 μM) and D2R antagonist Sulpiride (5 μM) on a single pulse (A), a single stimuli burst (B) and their ratio (C) in slices without tonic activity (*,**,**** - p < 0.05, 0.005 and 0.0001 by one-way ANOVA; n = 25 Ctrl, 14 Sulpiride and 7 DHβE). **(D)** Effect of artificial tonic activity on DA release evoked by a single burst (* - p<0.05 by two- way ANOVA; n = 46 (0 Hz), 25 (1 Hz) and 8 (2 Hz) for Ctrl; 10, 6 and 4 for DHβE; 13, 7 and 5 for Sulpiride). **(E-F)** Changes in the amplitude of DA release evoked by three sequential burst stimuli in the presence of DHβE (E) and Sulpiride (F; *,*** - p<0.05 or 0.001 by two-way ANOVA).

### 2.3. Role of D2R

Modulation of dopamine release within the striatum originates from a dynamic interplay between dopamine receptors and a local striatal circuitry. D2 auto-receptors expressed on the axons of DA neurons provide feedback control of striatal DA release via activation of potassium GIRK channels that hyperpolarize the cells ^18,38^, while D2 receptors on ChIs regulate ACh and DA release via postsynaptic mechanisms ^39,40^.

Acute treatment of striatal slice with D2R antagonist sulpiride (5 μM) had no effect on DA released by a single pulse stimulus, but increased the amount of DA overflow following burst stimulation (**Figure 3A-C**), consistent with previous reports ^18,19^. Tonic slice stimuli in the presence of sulpiride caused further decrease in DA released by a single burst (**Figure 3D**), consistent with a disinhibition of dopaminergic axons that produced larger depletion of the RRP of dopaminergic vesicles. Furthermore, blockade of D2R combined with tonic stimulation caused more DA to be released in consecutive bursts, leading to a facilitation of neurotransmitter release (**Figure 3F, Table 1**).

**Table 1.** Summary of changes in DA and ACh levels during tonic slice stimulation and nAChR blockade.

| Treatment | Tonic activity | Single pulse<br>*Figs 2C&3A* | Single burst<br>*Figs 3B&D* | Multiple bursts<br>*Figs 3E&F* |
| --- | --- | --- | --- | --- |
| Ctrl | - |  |  | depression |
| DHβE | - | ↓ | ↑↑ | depression |
| Sulpiride | - | ↔ | ↑ | depression |
| Ctrl | + | n.d. | ↓ | no change |
| DHβE | + | n.d. | ↓ | no change |
| Sulpiride | + | n.d. | ↓↓ | facilitation |
| *In vivo* | + | n.d. |  | facilitation |

n.d. is 'not detected'; ↔ indicated no difference relative to control.

## 3. DISCUSSION

In this study, we introduce a simple experimental approach that can be employed to study the role of tonic activity in modulating phasic DA release in acute striatal slice preparations. While the highest tested tonic frequency (5 Hz) caused complete cessation of DA neurotransmission (**Figure 2D**), likely due to the depletion of vesicular transmitter pools, at lower frequencies we uncovered several key characteristics of DA release that have not been previously reported in *ex vivo* preparations (**Table 1**).

First, tonic slice conditioning greatly diminished DA release following a single pulse stimulation, accompanied by a much smaller decrease in phasic neurotransmitter release, thus leading to an enhanced burst-to-single stimulus ratio. Such ‘high frequency filtering’ of DA neurotransmission, also observed *in vivo*, would lead to enhanced signal/noise during phasic DA release in response to environmental cues critical for learning ^41^. Interestingly, similar alteration in DA release kinetics has been reported in acute striatal slices treated with nAChR blockers ^12,20,33^ (**Figure 3C**), suggesting that DA and ACh tones have opposite effects on the signal/noise of DA neurotransmission. Nicotinic receptors are ligand-gated non-selective cation channels that potently modulate DA release in the striatum ^35,42,43^ and their activation on DA neurons drives depolarization and local action potentials ^44^. It is reported that tonic ACh facilitates DA release at lower firing frequencies but has no effect or causes depression of dopamine release at higher frequencies (>2 Hz) ^33,36,43,45^. The similarity between the effects of nAChR antagonists and the presence of tonic slice stimuli that both decrease single pulse but not burst DA release, may be explained by the desensitization of nAChR ^33^. During local tonic electrical stimulation of the slice, basal ACh concentration likely also increases, leading to nAChR desensitization. Since recovery from desensitization of nAChR may take up to one minute ^46^, at the time of the burst stimulus (10 sec after the end of the tonic train) the effect of the tonic conditioning would be similar to nAChR pharmacological blockade (**Table 1**), although the contribution of this and other mechanisms requires further investigation.

Another mechanism that exerts control over striatal DA neurotransmission is via activation of D2R expressed on both dopaminergic axons and ChI cell bodies ^19,39,47^. In acute striatal slices, tonically released DA is expected to bind D2R and induce chronic depression of DA axons and ChI, thus decreasing the probability of DA release. Conversely, D2R blockade not only allows more DA to be released from slices without tonic activity (**Figure 3A**), but also causes higher apparent synaptic vesicle depletion after tonic activity (**Figure 3D**).

As long reported, in naïve slices, sequential bursts of stimuli cause a profound depression of DA release ^17,48^, but we now show that tonic stimulation of the slice leads to less dependence of released DA on the burst number (**Figure 2G**). Furthermore, when tonic slice stimulation was combined with the pharmacological blockade of D2R, repeated stimulus bursts *facilitated* DA release, a response that has not been previously observed *ex vivo* (**Figure 3F**). A possible explanation for this phenomenon is that DA released from different vesicle pools possesses different calcium sensitivities. For example, if release from the RRP has higher Ca^2+^ affinity (i.e., can be triggered at lower cytosolic Ca^2+^ concentrations) than release from the recycling and the reserve pools, then depletion of the RRP during tonic activity would reveal synaptic facilitation due to Ca^2+^ buildup during high-frequency bursts. Consistently, we find that burst-to-burst depression of DA release was inversely proportional to tonic frequency, while release facilitation occurred in slices treated with a D2R antagonist, which as discussed above, drives still stronger depletion of the RRP. These phenomena can be different between dorsal and ventral striatum ^9,19^ and may further depend on muscarinic ACh receptors ^37^, GABA receptors ^49^ and other receptor-mediated mechanisms.

While the experimental system we introduce is relatively simple, it features many parameters that can be varied depending on the aims of the study, including the intensity and the frequency of tonic and phasic stimuli, the duration of tonic stimulation, and the delay before the burst stimulation and between the bursts. Furthermore, targeted optogenetic stimulation of DA axons and/or ChI cell bodies, as well as assessment of basal DA and ACh levels with electrochemical or optical techniques promise to further decipher the roles of tonic dopaminergic and cholinergic activities in modulating phasic DA neurotransmission.

In summary, we present proof of principle study that demonstrates the critical role of tonic firing in shaping dopamine release dynamics and highlights the importance of studying neurotransmitter modulation in physiologically relevant settings. Bridging the gap between *ex vivo* and *in vivo* data will pave the way for understanding the mechanisms underlying dopamine signaling and its implications for neurological disorders and pharmacological interventions.

## 4. MATERIALS AND METHODS

### 4.1. Experimental model and reagents

Male C57Bl6/J mice (3-6 months, The Jackson Laboratory, Bar Harbor, Maine) were used for both *in vivo* and *ex vivo* experiments. Animals were maintained according to the National Institutes of Health guidelines in Association for Assessment and Accreditation of Laboratory Animal Care (AAALAC) accredited facilities on a 12:12 hour light/dark cycle with food and water *ad libitum*. All experimental procedures were approved by the Columbia University Institutional Animal Care and Use Committee and followed guidelines established in the NIH Guide for the Care and Use of Laboratory Animals. Sulpiride (Cat#: 0894) and dihydro-β-erythroidine hydrobromide (Cat#: 2349) were from Tocris Bioscience.

### 4.2. *Ex vivo* striatal slices

Animals were euthanized by cervical dislocation, the brain was removed and placed in ice-cold sucrose cutting solution, containing (in mM): 10 NaCl, 2.5 KCl, 25 NaHCO3, 0.5 CaCl2, 7 MgCl2, 1.25 NaH2PO4, 180 sucrose and 10 glucose, oxygenated with 95% O2/5% CO2 to pH 7.4. Coronal slices (250 μm) that included the striatum were collected and allowed to rest at 34°C for 30 min in artificial cerebrospinal fluid (ACSF; in mM): 125 NaCl, 2.5 KCl, 25 NaHCO3, 1.5 CaCl2, 1 MgCl2, 1.25 NaH2PO4 and 10 glucose, oxygenated with 95% O2/5% CO2 to pH 7.4.

### 4.3. Fast-scan cyclic voltammetry (FSCV) *ex vivo*

FSCV recordings of evoked DA release by FSCV were performed as detailed previously ^36^. Striatal slices were incubated in oxygenated ACSF at room temperature for 30 min and transferred to a recording chamber with ACSF perfused at 2 mL/min and 34°C. A carbon fiber electrode was placed in the dorsolateral striatum ∼50 μm into the slice. Triangular voltage ramps from a holding potential of - 450 mV to +800 mV over 8.5 ms (ramp of 294 V/s) were applied to the carbon fiber electrode every 100 ms. Current was recorded with an Axopatch 200B amplifier (Molecular Devices, Foster City, CA) filtered with a 10-kHz low-pass Bessel filter and digitized at 25 kHz (ITC-18 board, InstruTECH, Great Neck, NY). FSCV data acquisition and analysis were controlled by custom-built routines in Igor Pro (WaveMetrics Inc., Lake Oswego, OR). Slices were stimulated with a sharpened bipolar concentric electrode (400 μm max outer diameter; Pt/Ir; WPI) placed ∼150 μm from the recording electrode, using an Iso-Flex stimulus isolator (AMPI) triggered by TTL pulses generated in the same custom-built Igor Pro routine and synchronized with FSCV pulses. The duration of each stimulus pulse was 0.2 ms. Carbon fiber electrodes were calibrated by quantifying background-subtracted voltammograms in standard solutions of DA in ACSF prepared fresh each recording day. Concentration traces were prepared in Igor Pro by collecting current over time at the peak DA oxidation potential normalized to the standard DA calibration factor.

### 4.4 FSCV recordings *in vivo*

We used previously published methods for *in vivo* FSCV recordings ^22^. Briefly, male mice were anesthetized with iso-flurane (induction 4%, maintenance 0.8–1.4%), placed on a heating pad, and head-fixed on a stereotaxic frame. Craniotomy was performed to insert a 22G bipolar stimulating electrode (P1 Technologies, Roanoke, VA) in the ventral midbrain to trigger electrical pulse trains and a carbon-fiber microelectrode (5 μm diameter, ∼150 μm length) in the dorsal striatum to measure evoked DA release. The depth of the stimulating electrode was adjusted between 4 and 4.5 mm for maximal DA release. FSCV recordings and generation of the stimulation patterns were performed as described above. The electrical pulse trains were delivered to the stimulating electrode at a constant current of 400 μA using an Iso-Flex stimulus isolator; duration of each stimulus pulse was 4 ms. The carbon-fiber microelectrodes were calibrated in artificial cerebrospinal fluid using known concentrations of DA.

### 4.5. Statistical Analysis

Data was acquired and analyzed using the Igor Pro software and imported into Microsoft Excel (Microsoft Corp., Redmond, WA). All statistical analysis was performed in GraphPad Prism (version 10; La Jolla., San Diego, CA). All bar graphs show the mean ± SEM. Data comparing two variables were analyzed with a two-way ANOVA, followed by Bonferroni post-hoc test. Data comparing one variable among more than two groups were analyzed with one-way ANOVA and Tukey’s post-hoc test.

### SAFETY

We have not found any unexpected, new, and/or significant hazards or risks associated with the reported work.

## ABBREVIATIONS

ChI: Cholinergic interneurons
DA: Dopamine
D2R: Dopamine D2 receptor
DHβE: Dihydro-β-erythroidine hydrobromide
FSCV: Fast-scan cyclic voltammetry
GABA: Gamma-aminobutyric acid
GIRK: G-protein-coupled inward rectifier potassium
MFB: Medial forebrain bundle
nAChR: Nicotinic acetylcholine receptors
RRP: Readily releasable pool
SNpc: Substantia nigra pars compacta

## AUTHOR CONTRIBUTIONS

SB, SJC and EAM conducted *ex vivo* FSCV experiments; EAM performed *in vivo* FSCV recordings in anesthetized mice. Data analysis and presentation was done by SB and EVM. All other authors contributed to experimental design, critical data analysis and writing of the manuscript.

## ACKNOWLEDGMENT

This research was supported by NIDA R0107418 and the JPB Foundation (DS), as well as the Fulbright Joint Supervision Program Award to SB. We thank Avery McGuirt for advice and training and Dr. Mark Wightman for critical discussions of the experimental approach.

